# ACE2-Targeting Monoclonal Antibody as Potent and Broad-Spectrum Coronavirus Blocker

**DOI:** 10.1101/2020.11.11.375972

**Authors:** Yuning Chen, Yanan Zhang, Renhong Yan, Guifeng Wang, Yuanyuan Zhang, Zherui Zhang, Yaning Li, Jianxia Ou, Wendi Chu, Zhijuan Liang, Yongmei Wang, Yili Chen, Ganjun Chen, Qi Wang, Qiang Zhou, Bo Zhang, Chunhe Wang

## Abstract

The evolution of coronaviruses, such as SARS-CoV-2, makes broad-spectrum coronavirus preventional or therapeutical strategies highly sought after. Here we report a human angiotensin-converting enzyme 2 (ACE2)-targeting monoclonal antibody, 3E8, blocked the S1-subunits and pseudo-typed virus constructs from multiple coronaviruses including SARS-CoV-2, SARS-CoV-2 mutant variants (SARS-CoV-2-D614G, B.1.1.7, B.1.351, B.1.617.1 and P.1), SARS-CoV and HCoV-NL63, without markedly affecting the physiological activities of ACE2 or causing severe toxicity in ACE2 “knock-in” mice. 3E8 also blocked live SARS-CoV-2 infection *in vitro* and in a prophylactic mouse model of COVID-19. Cryo-EM and “alanine walk” studies revealed the key binding residues on ACE2 interacting with the CDR3 domain of 3E8 heavy chain. Although full evaluation of safety in non-human primates is necessary before clinical development of 3E8, we provided a potentially potent and “broad-spectrum” management strategy against all coronaviruses that utilize ACE2 as entry receptors and disclosed an anti-coronavirus epitope on human ACE2.

## Introduction

In the last 20 years, coronaviruses have caused three major transmissible disease outbreaks in human, including severe acute respiratory syndrome (SARS) ^1^, Middle East respiratory syndrome (MERS) ^2^ and coronavirus disease 2019 (COVID-19) ^3,4^.

One of the challenges to control coronaviruses is that they evolve constantly, even though slower than HIV and influenza^5^. Analyses of over 28,000 gene sequences of SARS-CoV-2 spike protein (S-protein) in May 2020 revealed a D614G amino acid substitution (SARS-CoV-2-D614G) that was rare before March 2020, but increased greatly in frequency as the pandemic spread worldwide, reaching over 74% of all published sequences by June 2020 ^6^ and 81% by May 2021 (GISAID). Evolution of coronaviruses renders them ability to evade virus-specific medications ^7,8^. Recently, the emergence of multiple mutant variants of SARS-CoV-2, including B.1.1.7 (UK), B.1.351 (South Africa), P.1 (Brazil) ^9^ and B.1.617 ^10^ (India) manifests such challenge. In fact, a monoclonal antibody against SARS-CoV-2, bamlanivimab, has been revoked Emergency Use Authorization for expected poor performance against variants currently popular in the US (FDA news). In theory, broad-spectrum coronavirus therapeutics can withstand viral mutations and be potentially utilized in future campaigns against different coronavirus outbreaks.

The key to developing broad-spectrum coronavirus therapeutics is to identify broad-spectrum anti-viral targets. Although RNA polymerase is a broad anti-RNA virus target, it suffers from low specificity and efficacy ^11,12^. By employing a multi-dimensional approach, Gordon et al. proposed a set of potential “pan” viral target for coronaviruses, but the druggability of these targets are yet to be evaluated ^13^. ACE2 fusion proteins can act as decoy receptors to trap SARS-CoV-2 ^14,15^, but the affinity and developability of these proteins are generally less than antibodies. Recently, Rappazzo et al. generated a set of monoclonal antibodies that bound to a large panel of coronaviruses, but their neutralizing abilities have not been tested yet ^16^.

The infection of SARS-CoV-2 is triggered by binding of their envelope spike glycoproteins (S-protein) to angiotensin-converting enzyme 2 (ACE2) molecules expressed on host cells ^17,18^. The S-protein consists of two subunits: 1) S1-subunit (also called S1-protein) at N-terminal, containing the receptor-binding domain (RBD) responsible for ACE2 binding; 2) S2-subunit at C-terminal responsible for membrane fusion ^18^. The RBD of SARS-CoV-2 has been heavily targeted by antibodies as well as small molecule approaches ^19–23^, but the RBD-targeting approaches are prone to drug resistance caused by viral evolution and are not broad-spectrum.

ACE2 is a type-I transmembrane glycoprotein that plays important roles in maintaining blood pressure homeostasis in the renin-angiotensin system ^24,25^. It is a shared receptor for multiple coronaviruses, such as SARS-CoV-2, SARS-CoV, HCoV-NL63^17,26,27^, bat coronavirus RaTG13^28^, pangolin coronavirus GX/P2V/2017 and GD/1/2019 ^29^. SARS-CoV, a close sibling of SARS-CoV-2 in the coronavirus family, was the culprit that caused SARS outbreak in 2003 ^3^, while HCoV-NL63 infects human much more frequently but causes only cold symptoms with moderate clinical impacts ^30^. Binding of coronavirus to ACE2 not only facilitates viral entry into the host cells, but also down-regulates ACE2 expression ^31,32^.

Previous results revealed that the RBD binding site of ACE2 does not overlap with its catalytic site ^33–35^, it is therefore hypothesized that targeting the RBD binding site on ACE2 with antibodies can block the entry of all ACE2-dependent coronaviruses, while sparing ACE2’s physiological activities. Such antibodies can be theoretically utilized in managing both current and future coronavirus outbreaks and tolerate viral mutations. By targeting ACE2, additionally, the antibody could be evaluated in HCoV-NL63 patients even when COVID-19 patients are no longer available for clinical trials.

To test the hypothesis, we generated a monoclonal antibody, namely 3E8, to target the RBD-binding site on ACE2. The therapeutic potentials and safety profiles of 3E8 were investigated and the key binding sites of 3E8 on human ACE2 molecule were revealed by cryo-EM and mutation studies to aid future drug discovery endeavor.

## Results

### Antibody generation by hybridoma technology

BALB/c mice were immunized with Fc-tagged human ACE2 protein and the sera were screened for binding to ACE2 (supplementary Fig. 1a) and blocking of SARS-CoV-2 S1-subunit and ACE2 interaction (supplementary Fig. 1b). Hybridoma cells were constructed and the supernatants were screened by the same assays mentioned above. Antibody 3E8 was screened out from a pool of neutralizing antibodies as the most efficacious blocker of S1-subunit binding to ACE2. The variable regions of the heavy (V_H_) and light (V_L_) chains were cloned into human IgG4 backbone, transiently expressed in HEK293F cells and purified (supplementary Fig. 1c). Protein qualities were examined by reduced and non-reduced gels (supplementary Fig. 1d).

### 3E8 binds human ACE2 with moderate affinity

We measured the binding affinity of 3E8 to His-tagged human ACE2 protein with ELISA and biolayer interferometry (BLI). The *EC_50_* value was 15.3 nM in ELISA (Fig. 1a) and apparent dissociation constant (KD) was 30.5 nM in BLI using dimeric ACE2 (residues 19-740) as the soluble analyte (Fig. 1b). It also bound to HEK293F cells ectopically overexpressing human ACE2 and to Vero E6 cells endogenously expressing human ACE2, as demonstrated by flow cytometry (supplementary Fig. 1e).

**Fig. 1.**
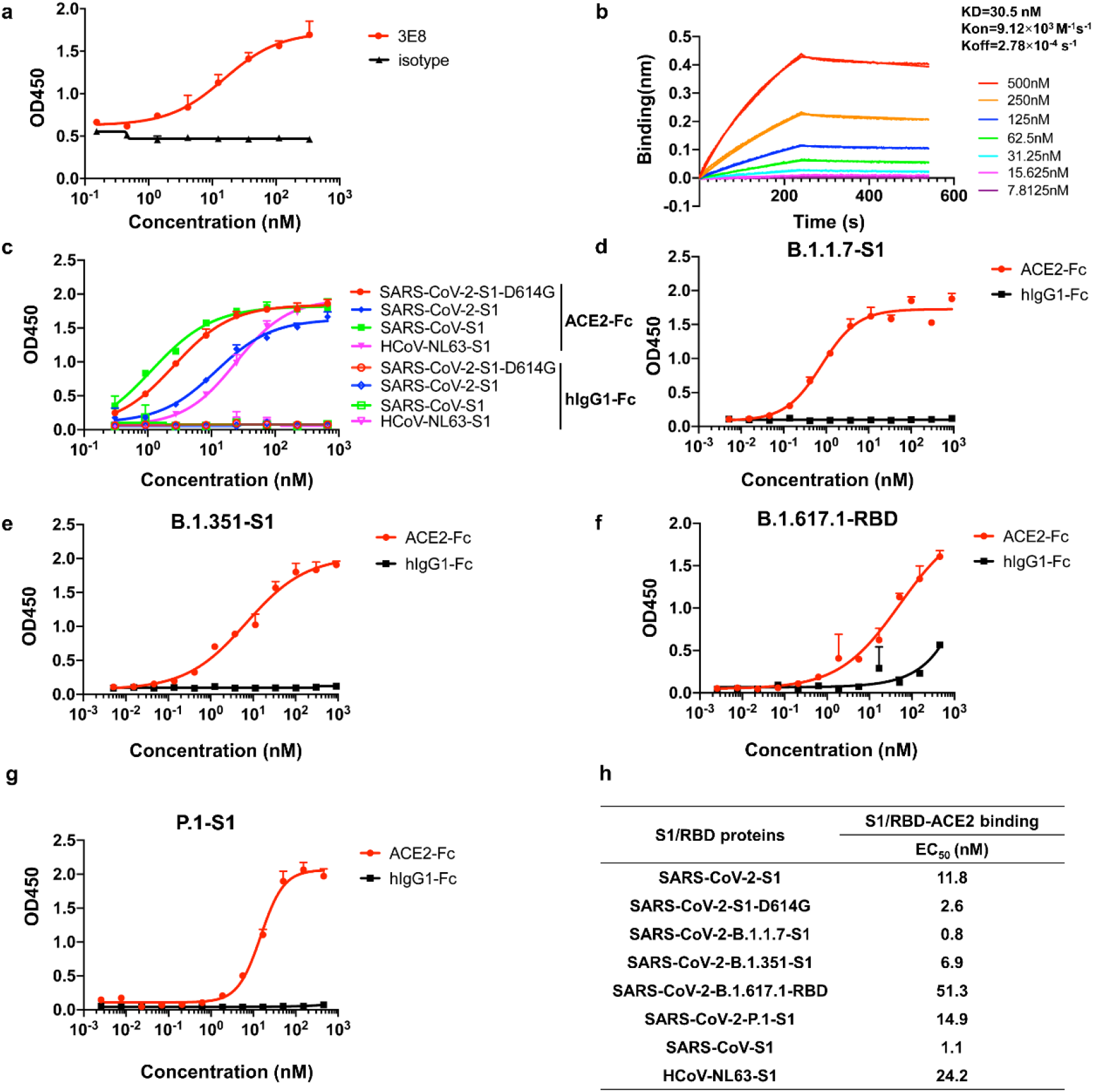
Monoclonal antibody 3E8 and recombinant S1-subunits or RBD from different coronaviruses (and SARS-CoV-2 mutant variants) bound to recombinant human ACE2 protein. **a** Binding of 3E8 to His-tagged recombinant human ACE2 protein as measured by ELISA. **b** Binding of 3E8 to His-tagged human ACE2 as measured by BLI. **c-g** Bindings of recombinant S1-subunits or RBD (in **f** only) from multiple coronaviruses and SARS-CoV-2 mutated variants to Fc-tagged recombinant human ACE2 protein as measured by ELISA. **h** The *EC_50_* values of recombinant S1-subunit bindings to human ACE2.

3E8 blocks the bindings of S1-subunits or RBD from multiple coronaviruses to ACE2 We investigated the abilities of 3E8 to block the ACE2 binding of S1-subunits or RBD from SARS-CoV-2, SARS-CoV-2-D614G, B.1.1.7, B.1.351, B.1.617.1, P.1, SARS-CoV and HCoV-NL63. These S1-subunits or RBD can all bind to Fc-tagged human ACE2 molecules (Fig. 1c-g), and the *EC_50_* values to Fc-tagged recombinant human ACE2 molecule were 11.8, 2.6, 0.8, 6.9, 51.3, 14.9, 1.1 and 24.2 nM, respectively (Fig. 1h). Incubation with 3E8 effectively blocked all S1-subunits or RBD binding to ACE2 (Fig. 2a-e) and the *IC_50_* values were 7.1, 13.8, 10.0, 3.7, 10.5, 9.3, 13.7 and 5.0 nM, respectively (Fig. 2f). Thus, 3E8 can broadly block the binding of S1-subunits or RBD from multiple coronaviruses, including the fast-spreading SARS-CoV-2 variants, to human ACE2 molecules.

**Fig. 2.**
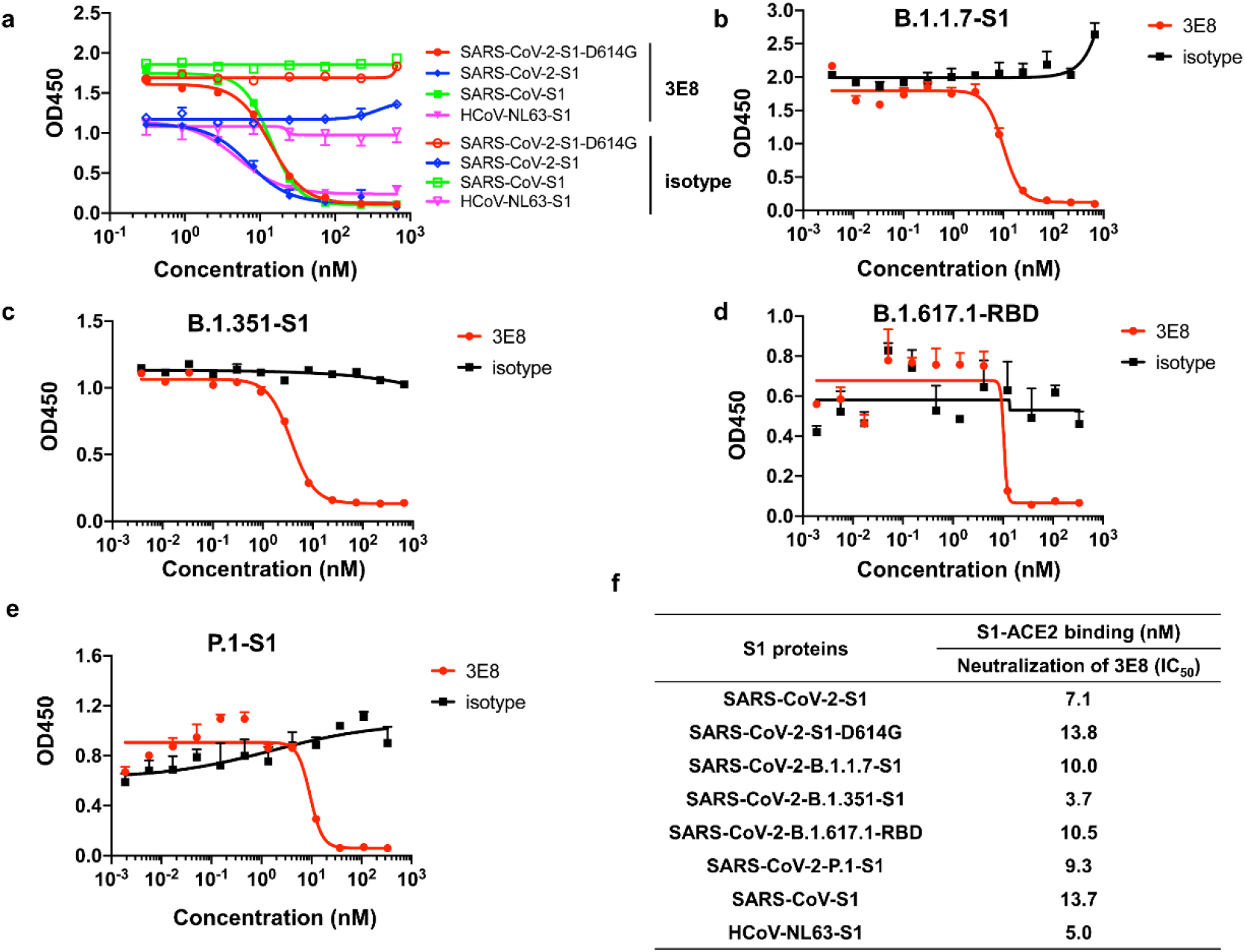
3E8 blocked the bindings of recombinant S1 or RBD from multiple coronaviruses and SARS-CoV-2 variants to Fc-tagged recombinant human ACE2 protein. **a-e** Bindings of His-tagged S1 or RBD (in **d** only) from different coronaviruses including SARS-CoV, HCoV-NL63, SARS-CoV-2 and emerging epidemic SARS-CoV-2 variants to recombinant human ACE2 protein were blocked by 3E8. **f** The *IC_50_* values of 3E8 in blocking S1 or RBD binding to human ACE2 protein.

### 3E8 abolishes the infectivity of multiple pseudo-typed coronaviruses

We next constructed pseudo-typed coronaviruses with full-length S-proteins from SARS-CoV-2, SARS-CoV-2-D614G, B.1.1.7, B.1.351, B.1.617.1, SARS-CoV and HCoV-NL63 (Fig. 3a-g). All pseudoviruses could infect HEK293F cells that ectopically express human ACE2, while SARS-CoV-2-D614G showed significantly enhanced infectivity when compared to the original SARS-CoV-2 (supplementary Fig. 2). Incubation with 3E8 fully abolished the infectivity of all pseudoviruses, with *IC_50_* values at 0.1, 0.1, 0.07, 0.3, 0.08, 0.2 and 1.1 nM, respectively (Fig. 3h). In comparison, B38, a SARS-CoV-2 RBD-targeting antibody currently under clinical development ^36^, could only suppress the infectivity of SARS-CoV-2, SARS-CoV-2-D614G, B.1.1.7 and B.1.617.1, but not B.1.351, SARS-CoV or HCoV-NL63. The suppression by 3E8 was not only broader, but also remarkably more efficacious and potent, as the *IC_50_* values of 3E8 was hundreds of folds improved when compared to that of B38 (Fig. 3h). ACE2-Fc fusion protein, a virus RBD-targeting molecule consisting the extracellular domain of human ACE2 and the Fc region of human IgG1, showed broad but moderate blocking ability on pseudoviruses. Our investigation indicated that 3E8 was potentially a powerful and broad-spectrum blocker on coronaviruses that are dependent on ACE2.

**Fig. 3.**
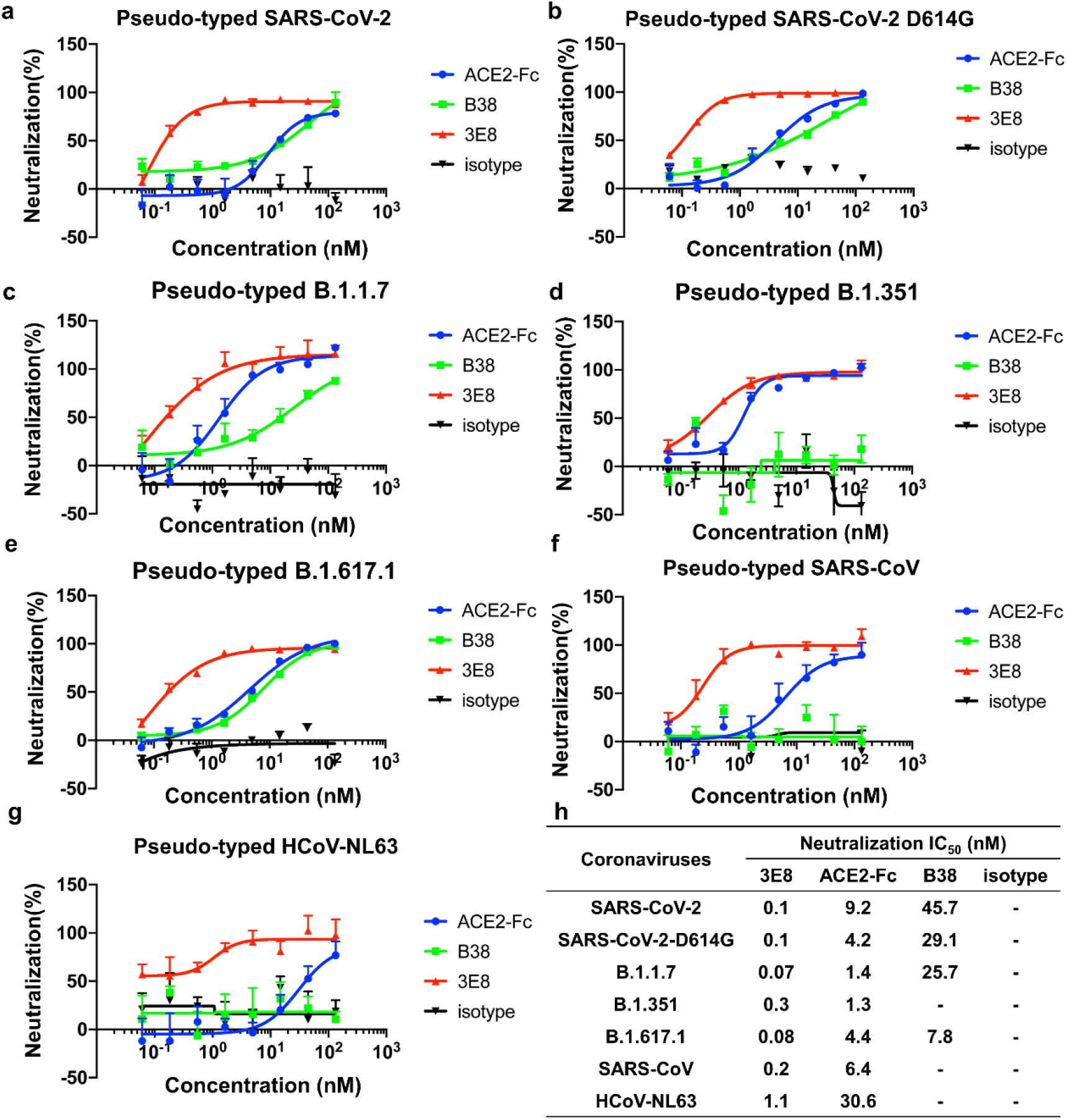
3E8 blocked the infections of ACE2-expressing cells by multiple pseudo-typed coronaviruses. ACE2-Fc and B38 were used as positive controls, and human IgG4 isotype was negative control. **a-g** 3E8 blocked infections of ACE2-overexpressing HEK293 cells by different pseudo-typed coronaviruses with Env-defective HIV-1 and full-length S-proteins from SARS-CoV-2, SARS-CoV-2-D614G, B.1.1.7, B.1.351, B.1.617.1, SARS-CoV and HCoV-NL63. **h** The *IC_50_* values of 3E8 in blocking pseudo-typed coronaviruses.

### 3E8 inhibits live SARS-CoV-2 infection of Vero E6 cells

Live virus study in a BSL-3 laboratory setting showed that incubation with 3E8 inhibited in a concentration-dependent manner the replication of SARS-CoV-2 in Vero E6 cells. The RBD-targeting B38 antibody, also inhibited SARS-CoV-2 replication, but was 60-fold less potent than 3E8, as suggested by the difference between their *IC_50_* values (2.3 vs. 0.04 nM), even though both completely abolished SARS-CoV-2 replication at high concentrations (Fig. 4a).

**Fig. 4.**
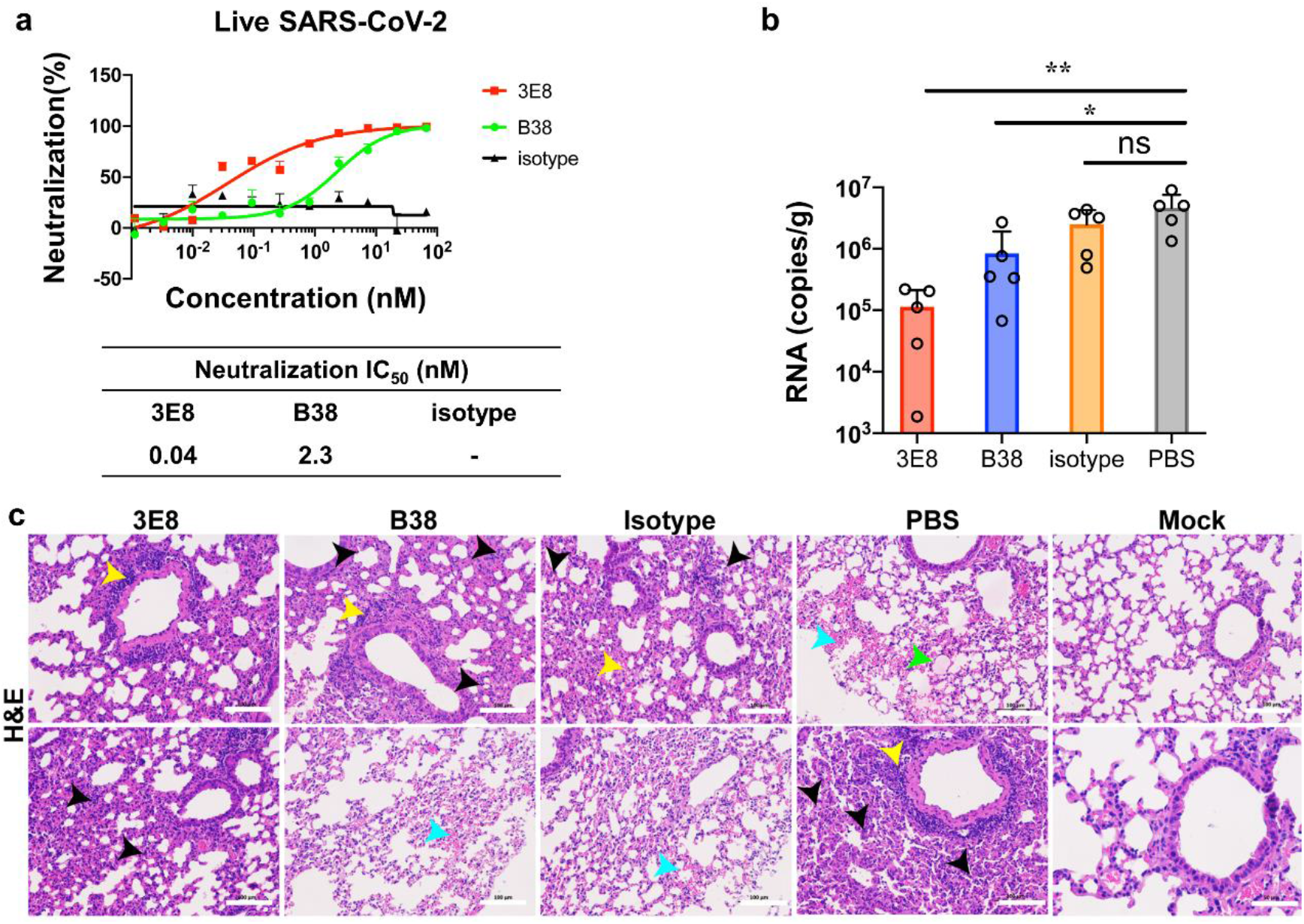
3E8 suppressed the infectivity of live SARS-CoV-2 in Vero E6 cells and a mouse model of COVID-19. **a** 3E8 suppressed the infection of Vero E6 cells by live SARS-CoV-2. **b** Application of 3E8 significantly reduced the viral RNA loads in the lungs of BALB/c mice ectopically expressing human ACE2 and inoculated with live SARS-CoV-2 virus. RBD-targeting monoclonal antibody B38 and isotype were used as positive and negative controls, respectively. **c** H&E staining of lung organ samples from different treatment groups. PBS- and isotype-treated mice developed serious interstitial pneumonia characterized with large area of alveolar septal thickening, large number of inflammatory cell infiltration (black arrow), even formed vascular cuff around blood vessels (yellow arrow), bleeding areas (blue arrow), and material exudates from the alveolar cavity (green arrow). B38-treated mice showed slightly-less histological pneumonia than the two groups. Only inflammatory cell infiltration (black arrow) and formed vascular cuff around blood vessels (yellow arrow) were observed in the lungs of 3E8-treated mice. The scale represents 100 μm. *: *p*<0.05; **: *p*<0.01.

### 3E8 blocks SARS-CoV-2 in a prophylaxis mouse model of COVID-19

More importantly, the neutralizing ability of 3E8 was validated in a prophylaxis mouse model of COVID-19. This model was generated by exogenous delivery of hACE2 with Venezuelan equine encephalitis replicon particles, VEEV-VRP-hACE2 ^37^. It was a non-lethal infection model that was suitable for measuring viral RNA loads and histological pathology of lungs. After hACE2 was intranasally delivered by VEEV, the antibodies were administered intraperitoneally at 10 mg/kg, and mice were challenged intranasally with 10^5^ PFU of live SARS-CoV-2 12 hours later. The viral RNA loads and tissue damages in lungs were examined 3 days post infection. Consistent with our *in vitro* results, application of 3E8 protected lungs from virus infection, as indicated by approximately 40-fold reduction in lung viral loads (Fig. 4b) and ameliorated tissue damages (Fig. 4c). In comparison, the viral loads in B38-treated mice were only about 5 times lower than that of control mice. Thus, application of 3E8 achieved significantly greater anti-viral effects than that of B38 in the COVID-19 mouse model we employed.

### 3E8 has no effects on ACE2’s catalytic activities or causes toxicity in “knock-in” mice

Since ACE2 plays important roles in maintaining blood pressure homeostasis in the renin-angiotensin system, we evaluated the safety risks of 3E8 application both *in vitro* and *in vivo*. Our studies with both recombinant ACE2 protein and Vero E6 cells suggested that 3E8 had no effects on ACE2’s catalytic activities even at 666.7 nM (Fig. 5a, b). Furthermore, incubation with 3E8 did not trigger a clear trend of ACE2 degradation in Vero E6 cells, as indicated by Western blot (Fig. 5c). Although 3E8 caused time-dependent internalization of ACE2, the levels of membrane-expressed ACE2 were stabilized after 24 h of incubation (Fig. 5d). In limited number of human ACE2 “knock-in” mice, which express human instead of mouse ACE2, injection of 3E8 did not induce noticeable changes in body weights or blood chemistry profiles (supplementary Fig. 3). In addition, there were no obvious differences in shape, size or pathological staining of major organs, including hearts, livers, kidneys, spleens and lungs of treated mice (supplementary Fig. 3).

**Fig. 5.**
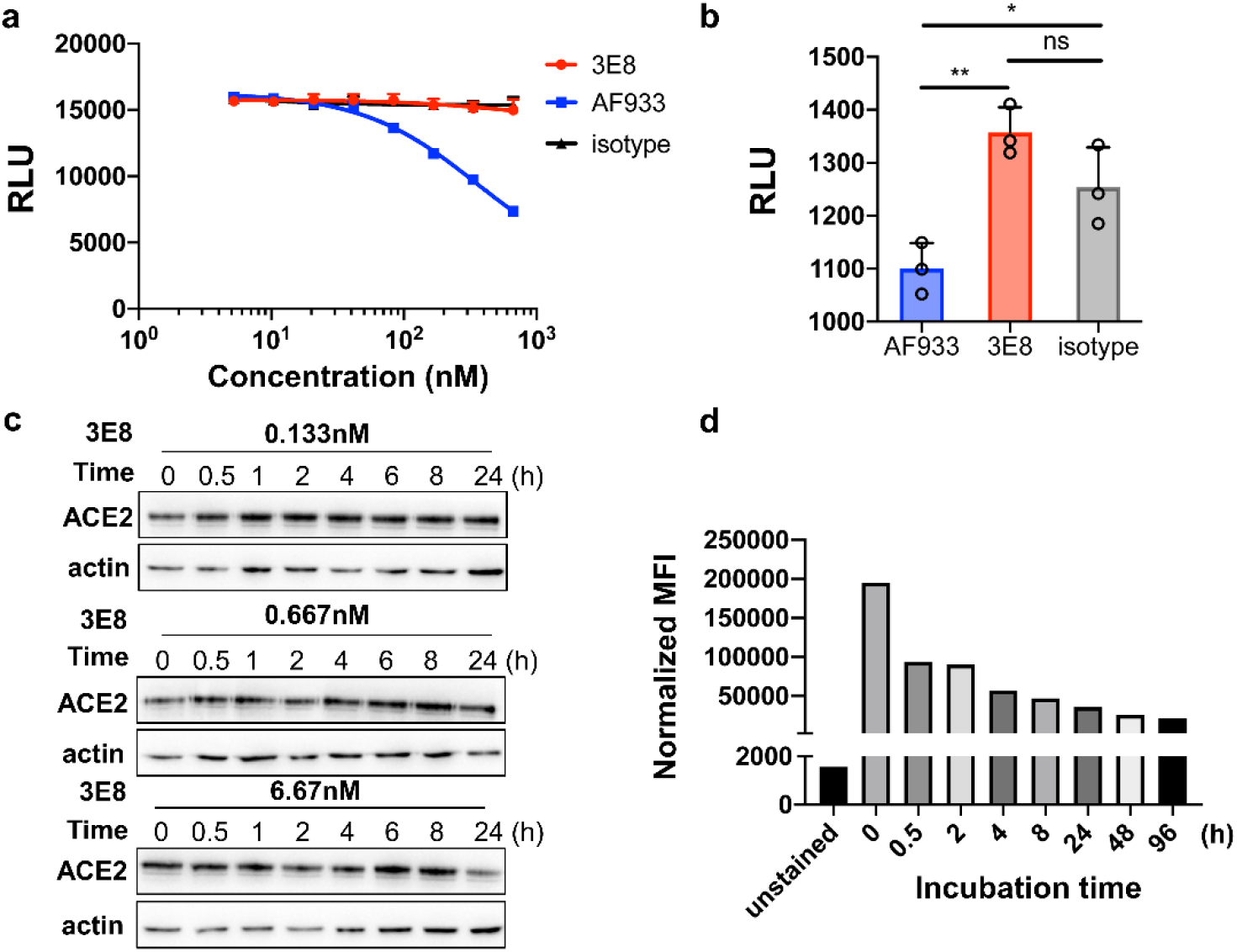
3E8 treatment had no effects on ACE2’s physiological functions. **a** 3E8 treatment had no effects on enzymatic activities of recombinant human ACE2 protein. AF933, an ACE2 polyclonal goat antibody, was used as positive control. hIgG1-Fc and isotype were used as negative controls. **b** 3E8 treatment showed no effects on enzymatic activities of ACE2 molecules endogenously expressed on the surface of Vero E6 cells. **c** Total expression of ACE2 by Vero E6 cells was not affected by 3E8 treatment at 0.133, 0.667 or 6.67 nM for 24 h. **d** The levels of membrane ACE2 expression on HEK293 cells were reduced by 3E8 treatment but were stabilized after 24 h. *: *p*<0.05; **: *p*<0.01.

### 3E8 binding epitope on ACE2 is determined by cryo-EM and “alanine walk” studies

To characterize the epitope recognized by 3E8 on ACE2, we solved the cryo-EM structure of the ACE2-B^0^ AT1 complex bound with 3E8 at an overall resolution of 3.2 Å (Fig. 6). Each ACE2 molecule in the complex is bound by a 3E8 molecule that extends from the complex like a wing (Fig. 6a). The heavy chain of 3E8 binds to the peptidase domain of ACE2 mainly through polar interactions between the complementarity-determining region (CDR) 2 and 3 of 3E8 and the N-terminal α1 helix of ACE2 (Fig. 6b). The loop between α2 and α3 of ACE2, referred to as Loop2-3, also contribute limited interactions with 3E8. The resolution at the interface was improved to 3.4 Å by applying focused refinement, supporting detailed analysis on the interactions between ACE2 and 3E8. The interface can be divided into two clusters. At cluster 1, the side chains of Asp103 and Arg104 of 3E8 are hydrogen (H) bonded with the main chain of Phe28 in α1 helix of ACE2 and the side chain of Tyr83 in Loop2-3 of ACE2, respectively (Fig. 6c). Meanwhile, the main chain atoms of Asp103 and Asp104 of 3E8 form H-bonds with the side chain of Gln24 of ACE2. At cluster 2, Tyr54 and Tyr102 of 3E8 interact with Lys31 of ACE2 through cation-π interactions, whereas Asn55 and Lys59 of 3E8 interact with His34 of ACE2 and Glu23 and Gln18 of ACE2, respectively, by forming H-bonds between side chains of these residues (Fig. 6d). Additionally, we performed “alanine walk” studies and identified Gln24 as the most critical amino acid residue that interact with the CDR3 of 3E8 heavy chain (Fig. 6e), consistent with the general concept that the CDR3 of heavy chain contributes the most to antigen recognition and binding. Although His34 was indicated by EM study to interact with the CDR2 of the heavy chain, mutation of it to alanine (Fig. 6e) or other amino acid residues (data not shown) failed to alter the binding of 3E8 to ACE2. Structural alignment of the 3E8/ACE2-B^0^ AT1 complex with the previously reported RBD/ACE2-B^0^ AT1 complex reveals clash between 3E8 and RBD of SARS-CoV-2 at the binding interface with ACE2 (Fig. 6f), providing an explanation for the results of competition assays. The binding site of 3E8, SARS-CoV-2, SARS-CoV and HCoV-NL63 on ACE2 were summarized ^38–40^ (Fig. 6g). Evolutionary tree of 3E8 binding site on ACE2 with different species was also provided (Fig. 6h), and phylogenetic diversities at position 23, 24, 31 and 34 were identified.

**Fig. 6.**
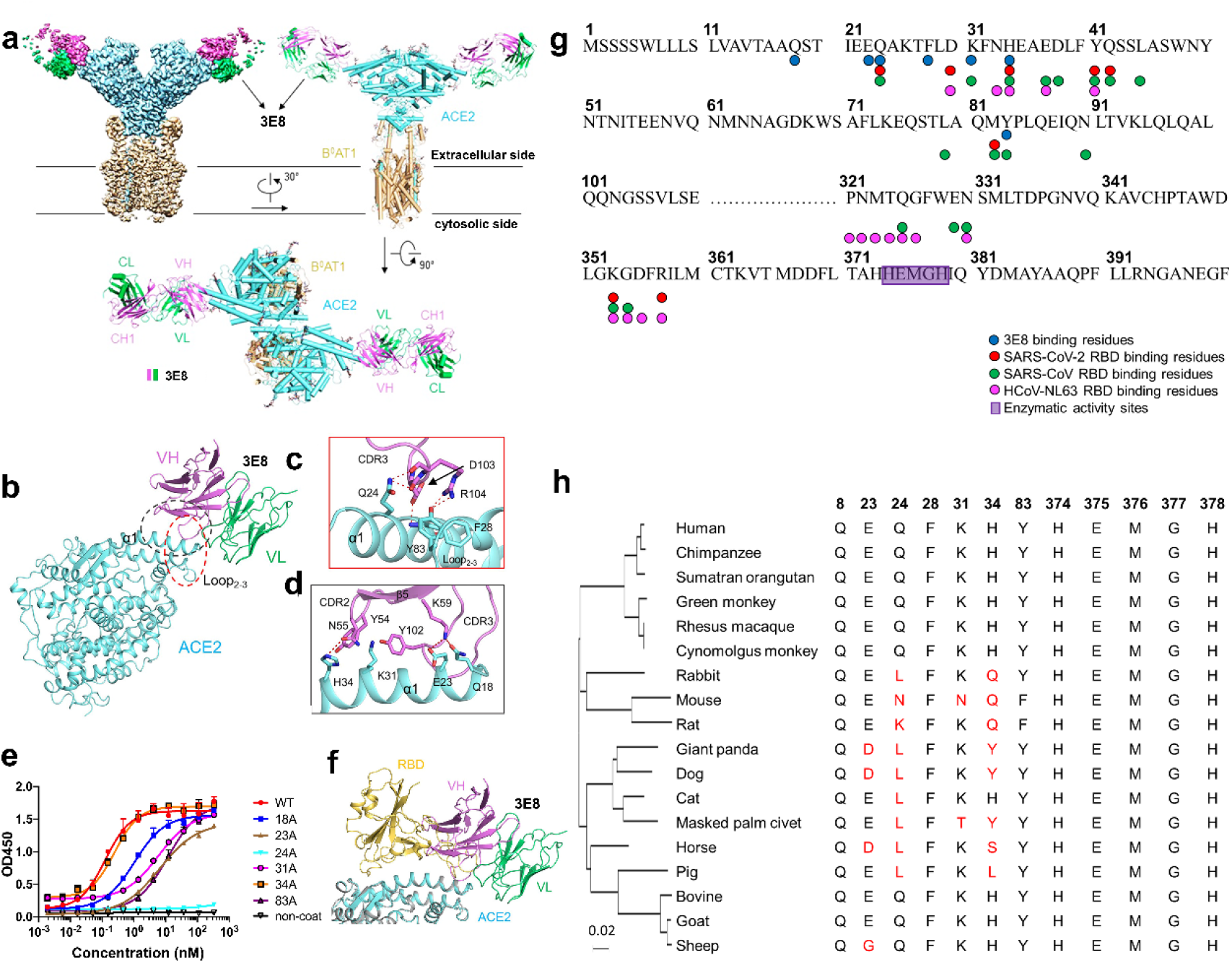
Cryo-EM structure of the 3E8/ACE2-B^0^ AT1 complex and “alanine walk” studies to solve the critical interactions between 3E8 and ACE2. Domain-colored cryo-EM map (**a**, upper left panel) and the side view (**a**, upper right panel) or the top view (**a**, lower panel) of the structure. The heavy and light chains of 3E8 are colored green and violet, respectively. The ACE2 and B^0^AT1 are colored cyan and wheat, respectively. **b** Binding interface between 3E8 and ACE2, which contains two clusters that are labeled with red and black dashed ellipses and detailed shown in **c** and **d**, respectively. H-bonds are indicated by red dashed lines. Q (Gln) 24 and H (His)34 on ACE2 were identified as critical amino acid residues interacting with the CDR3 and CDR2 of heavy chain of 3E8, respectively. **e** “Alanine walk” identified Q24 on ACE2 as the most critical amino acid residue for 3E8 binding. **f** Structural alignment of the 3E8/ACE2-B^0^AT1 complex and the RBD/ACE2-B^0^AT1 complex (PDB ID: 6M17) shows clash between 3E8 and RBD of the SARS-CoV-2 S protein. **g** Summary of binding site of multiple coronaviruses on human ACE2. **h** Evolutionary tree of 3E8 binding site on ACE2 with different species.

## Discussion

To block the entry of coronavirus, either virus- or ACE2-directed strategies can be taken. Targeting viruses directly is theoretically safer but vulnerable to viral evolution. By targeting ACE2 with an RBD-blocking antibody, we achieved broader and more effective suppression against ACE2-dependent coronaviruses without causing severe side effects. Furthermore, we revealed a broad-spectrum anti-coronavirus epitope on ACE2.

The mechanism by which 3E8 is more potent and efficacious than RBD-targeting antibody B38 is not yet fully understood. Limited by the sample size and evaluation using only a prophylactic regimen in a non-lethal animal model, it is premature to conclude that targeting ACE2 is superior to targeting viral RBD in potency or efficacy. B38 is one of the early anti-SARS-CoV-2 antibodies isolated from COVID-19 patients and due to the urgent nature, it was not well engineered with respect to affinity and developability. More head-to-head studies with more ACE2- and RBD-targeting molecules are necessary before drawing any conclusion.

ACE2-Fc (or called ACE2-Ig) fusion protein molecules may act as a “decoy” to interfere coronaviruses from binding to endogenous ACE2 molecules (Fig. 2 and Fig. 3). Although ACE2-Fc molecules are broad-spectrum in theory, their binding affinity (to RBDs), specificity and developability are usually lower than antibodies. ACE2-Fc was included in our studies as a control and moderate efficacy was observed *in vitro*. Thus, ACE2-neutralizing antibody appears to be a more favorable approach than ACE2-Fc fusion protein.

“Cocktails” or combination therapies have been currently explored in treating COVID-19 ^19^. A combination of 3E8 with antibodies recognizing different epitopes (e.g., RBD, NTD and/or glycan) on the viral surface seems a viable option and could be explored in clinic for better efficacy.

It is not surprising that no severe side effects or toxicity of 3E8 were observed *in vitro* or in human ACE2 “knock-in” mice. *In vitro*, 3E8 did not affect the catalytic activities of ACE2 or trigger significant ACE2 down-regulation. Even though ACE2 internalization was overserved, the levels of membrane ACE2 expression were stabilized after 24 h. It is possible that the ACE2 molecules remaining on the membrane are sufficient to maintain the physiological functions of ACE2-expressing cells. Previous studies showed that ACE2 “knockout” mice were viable and healthy in general, even though the contractile dysfunction was found ^41^, indicating that ACE2 is not crucial to the survival of animals. Due to limited animal availability, the conclusion from human ACE2 “knock-in” mice should not be overinterpreted. We plan to repeat this study when more animals are commercially available. Moreover, key signs of cardiovascular health, such as pulse pressure and heartbeat rate, cannot be conveniently measured in mice. Thus, the side effects and toxicities of 3E8 should be carefully evaluated in non-human primates before moving to the clinic.

A few broad-spectrum anti-coronavirus antibody or virus decoy receptor strategies have been disclosed ^14,15,16^. Among them, Rappazzo et al. reported an RBD-targeting antibody with exceptional breadth against distantly related SARS viruses ^16^, even though some highly transmissible variants weren’t explicitly tested since they were not well described in January 2021 when Rappazzo’s manuscript was accepted for publication. The antibody performed well in a phase 1 trial and is currently in a phase 2/3 trial, exemplifying the power of broad-spectrum antibody therapies. Nevertheless, current coronavirus-targeting antibodies focus mainly on highly conserved regions of RBD, such as S309 ^20^, 47D11 ^21^, D405, G502, G504 and Y50 ^16^. The epitope of 3E8 binding on ACE2 is only partially overlapping with that of RBD domain, but blocked virus infections with remarkable efficiency, demonstrating the extraordinary power of ACE2 targeting strategy. Previously, neutralizing antibodies (4A8) ^42^ and 89C8-ACE2 ^43^ were isolated from convalescent COVID-19 patients with binding on the N-terminal domain (NTD) of the SARS-CoV-2 S-protein, but not the RBD. Our results highlighted again the importance of epitope outside or on the verge of RBD/ACE2 interface, and would facilitate future endeavor searching for broad-spectrum anti-coronavirus approaches.

Overall, we presented evidence that 3E8 was a promising therapeutic candidate for coronavirus pandemic and believe that it represents a significant conceptual advance in fighting COVID-19, which keeps evolving, and may open the door for more ACE2-targeting drug discovery and development.

## Materials and methods

### Spike mutations on SARS-CoV-2 variants

B.1.1.7: H69 deletion, V70 deletion, Y144deletion, N501Y, A570D, D614G, P681H, T716I, S982A, D1118H; B.1.351: D80A, D215G, A242-244 deletion, K417N, E484K, N501Y, D614G, A701V; B.1.617.1: T95I, G142D, E154K, L452R, E484Q, D614G, P681R, Q1071H; P.1: L18F, T20N, P26S, D138Y, R190S, K417T, E484K, N501Y, D614G, H655Y, T1027I, V1176F.

### Biolayer interferometry (BLI)

Binding affinities were measured by BLI using Fortebio Octet Red 96. For affinity measurement, 10 μg/ml of 3E8 was captured by protein A biosensor and incubated with different concentrations of hACE2-his protein. The baseline was established by PBS with 0.05% tween-20 for 60 s. The association was set at 240 s and the dissociation periods was set at 300 s. The mean Kon, Koff and apparent KD values of binding affinities were calculated from all binding curves based on their global fit to 1:1.

### Neutralization ELISA

S1-proteins (2 μg/ml) were coated onto plates at 4°C overnight. Serial dilutions of 3E8 or isotype were pre-incubated with 5 μg/ml of ACE2-Fc for 30 min at room temperature. Then, the mixture was added into the coated plate wells and incubated for 1h. The bound ACE2-Fc was detected by HRP-conjugated goat anti-human IgG and developing substrate.

### Pseudo-typed virus neutralization assay

Pseudo-typed SARS-CoV-2-D614G, SARS-CoV-2, SARS-CoV, HCoV-NL63, B.1.1.7, B.1.351 and B.1.617.1 were constructed by co-transfection of two plasmids, one expressing Env-defective HIV-1 with luciferase reporter (pNL4-3.luc.RE)^44^ and the other expressing the full-length S-protein of SARS-CoV-2-D614G, SARS-CoV-2, SARS-CoV, HCoV-NL63, B.1.1.7, B.1.351, or B.1.617.1 into HEK293T cells. The supernatant containing virus particles was harvested 48 h post-transfection followed by 0.45 μm filtration. HEK293F/ACE2/EGFP cells were pre-seeded with 1.2×10^4^ cells per well in a 96-well plate. The confluent cells were incubated with 50 μl of serially-diluted antibodies or ACE2-Fc for 1h at 37°C followed by addition of various pseudoviruses of the same volume. 100 μl of DMEM with 10% FBS was added as negative control. 100 μl of pseudovirus and DMEM mixed at ratio 1:1 was used as positive control. After 24 h, medium was changed and the cells were incubated for another 48 h. The relative light units (RLUs) of luminescence were measured by Firefly Luciferase Reporter Assay Kit (Meilunbio). Neutralization (%) = [1- (RLU_samples_ - RLU_negtive control_) / (RLU_positive control_ - RLU_negtive control_)] × 100%. The *IC_50_* values were calculated by non-linear.

### Live SARS-CoV-2 suppression *in vitro*

Vero E6 (ATCC® CRL-1586™) cells were trypsinized and pre-seeded into 24-well plates in duplicate with 1 × 10^5^ cells/well in DMEM containing 10% FBS (100 U/mL of penicillin and 100 μg/ml of streptomycin) at 37°C with 5% CO_2_ one day before. After confluent, antibodies of 3-fold serially diluted were added, and Vero E6 cells were infected with live SARS-CoV-2 (IVCAS 6.7512) virus at a multiplicity of infection (MOI) of 0.01. After 24h incubation, the culture supernatants were collected and viral RNA was quantified via qRT-PCR using Luna® Universal Probe One-Step RT-PCR Kit (E3006) on CFX96 Touch™ Real-Time PCR Detection System (Bio Rad). Primers used were as follows: RBD-qF1: 5’-caatggtttaacaggcacagg-3’, RBD-qR1: 5’-ctcaagtgtctgtggatcacg-3’, Probe: acagcatcagtagtgtcagcaatgtctc. IC_50_ was fitted and calculated by GraphPad Prism 8. Data represents as mean ± SD of two replicates from one representative experiment, and the experiment was repeated for 3 times.

### COVID-19 mouse model and disease suppression

BALB/c mice were purchased from Wuhan Institute of Biological Products Co. Ltd. and cared in accordance with the recommendations of National Institutes of Health Guidelines for the Care and Use of Experimental Animals. All the animal studies were conducted in biosafety level 3 (BSL-3) facility at Wuhan Institute of Virology under a protocol approved by the Laboratory Animal Ethics Committee of Wuhan Institute of Virology, Chinese Academy of Sciences (Permit number: WIVA26201701).

A mouse model recently established by VEEV-VRP delivery of ACE2 for SARS-CoV-2 infection ^37^ was used to evaluate the efficacy of 3E8 *in vivo*. Four groups of six- to eight-week-old female BALB/c mice (n=5 per group) were first intranasally infected with 10^6^ FFU VRP-ACE2 per mouse in a total volume of 80 μl after anesthetization with Avertin (250 mg/kg). The mice from different groups were treated with 3E8, B38 or isotype at a dose of 10 mg/kg 12 h later via intraperitoneal injection. After another 12 h, all mice were intranasally infected with 10^5^ PFU SARS-CoV-2 in a total volume of 50 μl. 3 days post infection of SARS-CoV-2, the lungs of mice were collected for viral RNA quantification and histological analysis. For RNA quantification, some lungs were homogenized in DMEM medium, and viral RNA was extracted using QIAamp viral RNA mini kit (52906, Qiagen) following the manufacturer’s protocol. qRT-PCR assay was performed using Luna® Universal Probe One-Step RT-PCR Kit (E3006). Primers and probe used were: RBD-qF1: 5’-caatggtttaacaggcacagg-3’, RBD-qR1: 5’-ctcaagtgtctgtggatcacg −3’, Probe: acagcatcagtagtgtcagcaatgtctc. For histological analysis, lung samples from mice were fixed with 4% paraformaldehyde, embedded in paraffin, sagittally sectioned at 4 μm thickness on a microtome, and mounted on APS-coated slides for H&E stain.

### ACE2 enzymatic activity assay

The catalytic activities of recombinant and endogenous ACE2 was detected according to a published protocol using fluorescent substrate, Mca-APK-Dnp (AnaSpec) ^45^. To determine the impact on enzyme activity, serial diluted antibodies were pre-incubated with 2.0 μg/ml of recombinant human ACE2 at room temperature for 1 h on a shaker. After incubation, the neutralization solution was 1:5 diluted with activity buffer ^45^ and then mixed with 50 μl/well of 200 mM substrate. The mixture was incubated at 37 °C for 20 min before the RFU of fluorescent signals were read on an Envision microplate reader (Perkin Elmer, Waltham, MA) with excitation wavelength set at 320 nm and emission wavelength set at 400 nm. To measure endogenous ACE2 enzymatic activity, Vero E6 cells were seeded in a 96-well plate at 1×10^5^ cells/well and cultured overnight. The cells were then incubated with 10 μg/ml of 3E8, AF933(R&D Systems, Minneapolis, MN) and isotype for 60 min at 37°C before mixed with 50 μl/well of activity buffer and 50 μl/well of substrate. The cells were incubated for 20 min at 37°C before transferred to black 96-well plate for fluorescence reading.

## Data availability statements

All data supporting the findings of this study are available from the corresponding author on reasonable request.

## Acknowledgements

We thank Dr. Hong Qiu (Shanghai Institute of Materia Medica) for providing the eukaryotic codon-optimized SARS-CoV-2 S-protein gene, James C. Wang (Shanghai American School) for assistance in ELISA and manuscript editing, Dr. Lu Lu (Fudan University) for providing pNL4-3.luc.RE, the cryo-EM facility and supercomputer center of Westlake University for providing cryo-EM and computing support.

This work was supported by the China National Major Scientific and Technological Special Project for “Significant New Drugs Innovation and Development” (2019ZX09732002-006), the Strategic Priority Research Program of the Chinese Academy of Sciences (CAS) (XDA12020223 and XDA12020330), the National Natural Science Foundation of China (81872785, 81673347, 31971123, 32022037, 81920108015 and 31930059), Shanghai Municipal Commission of Science and Technology of China (17431904400 and 19YF1457400), Institutes for Drug Discovery and Development, Chinese Academy of Sciences (CASIMM0120202008, CASIMM0120202007), the National Key R&D Program (2020YFA0509303), Major Scientific and Technological Special Project of Zhongshan City (191022172638719, 210205143867019), the Key R&D Program of Zhejiang Province (2020C04001), the SARS-CoV-2 emergency project of the Science and Technology Department of Zhejiang Province (2020C03129), the Leading Innovative and Entrepreneur Team Introduction Program of Hangzhou, Westlake Education Foundation and Tencent Foundation.

## Conflict of interests

Yili Chen, Ganjun Chen and Chunhe Wang are employed by Dartsbio Pharmaceuticals.

## Contributions

Yuning Chen, Guifeng Wang, Jianxia Ou, Wendi Chu, Zhijuan Liang, Yongmei Wang and Qi Wang in Shanghai institute of Materia Medica constructed the hybridoma cells, screened the antibody and assayed the binding and neutralizing activity against ACE2 and pseudo-typed viruses; Yanan Zhang, Zherui Zhang and Bo Zhang in Wuhan Institute of Virology conducted the anti-live virus activities (both *in vitro* and in mouse models) and drafted the respective setion of the manuscript; Renhong Yan, Yuanyuan Zhang and Qiang Zhou in Westlake University conducted the cryo-EM experiments and drafted the respective setion of the manuscript; Yili Chen and Ganjun Chen in Dartsbio Pharmaceuticals conducted some of the molecular and cellular experiments; Yaning Li in Tsinghua University conducted the 3E8-ACE2 structural analysis experiments. Chunhe Wang designed the experiments and drafted the manuscript.

## Supplementary Materials for

### Materials and Methods

#### Protein expression and purification

The His-tagged S1 proteins of SARS-CoV, HCoV-NL63, SARS-CoV-2, SARS-CoV-2 D614G, B.1.17, B.1.351, P.1 and the His-tagged RBD protein of B.1.617.1 were purchased from Sino Biological (Beijing, China). ACE2-his, ACE2-Fc and 3E8 were prepared as below: The DNA fragments encoding extracellular ACE2 domain (residues 19-740) from plasmid encoding full-length ACE2 (Sino Biological) were subcloned into the mammalian expression vectors pTT5 and pINFUSE (Vivogen, San Diego, CA) with C-terminal 6×His or hIgG1-Fc tags. The codon-optimized variable regions of the heavy and light chain of 3E8 were cloned into expression vectors containing human IgG4 constant regions. Purified plasmids were transfected into HEK293F cells (Shanghai Cell Line Bank, China) by polyethylenimine (Polysciences, Warrington, PA). Cells were then cultured in suspension in CD medium. After 5 days of culture, the supernatant was collected and purified by Ni-NTA or protein A chromatography. Size exclusion chromatography column (SEC) was used to examine the purity of proteins.

#### Binding ELISA

96-well Immuno-plates (Greiner) were coated with 2.0 μg/ml of purified recombinant 6×His-tagged human ACE2 protein at 4°C overnight. After blocking at room temperature for 1 h with 1% casein (Thermo Fisher), plates were washed with PBS containing 0.05% Tween-20, and serial dilutions of 3E8 were added for 1 h incubation. After washing, goat anti-human IgG conjugated with HRP (AB Clonal Technology, 1:2000 dilution) was added and incubated for 1 h, then TMB substrate (Thermo Fisher Scientific) and 2 M of H_2_SO_4_ were added, and OD_450_ was detected with SpectraMax M5e (Molecular Devices) microplate reader. To measure the binding affinity of S1 proteins to ACE2, 2 μg/ml of various S1 proteins were coated onto plates followed by addition of gradient diluted ACE2-Fc, and goat anti-human IgG conjugated with HRP was used for detection.

#### Flow Cytometry

Vero E6 and HEK293/ACE2/EGFP cells were harvested and aliquoted into FACS tubes at 5×10^5^ cells/tube. The cells were washed with cold staining buffer (PBS+0.1% BSA+0.04% Na3N) and then resuspended in 100 μl of 3E8 at different concentrations. The cells were kept at 4°C in the dark for 1 h on a shaker before washed twice. The cells were resuspended in 100 μL of staining buffer containing PE-goat-anti-human IgG (Biolegend) at 4 μg/ml for 30 min. The cells were washed twice and resuspended in 200 μl of staining buffer for flow cytometry analysis.

#### Western Blot

Vero E6 cells were incubated at 37°C with different concentrations of 3E8 in DMEM with 10% FBS and lysed with commercial cell lysis buffer (Beyotime, Shanghai, China) at different time points. Western blot analysis for ACE2 protein in whole cell lysate was carried out using rabbit anti-ACE2 pAb (1:500, Sino biological) and goat anti-rabbit conjugated with HRP (1:2000, Abclonal, Wuhan, China) using a standard Western blot protocol.

#### Cryo-EM sample preparation

The ACE2-B^0^AT1 complex was mixed with 3E8 at a molar ratio of 1:1.5 at 4 °C for 1hr before applied to the grids. Aliquots (3.3 μl) of the protein complex were placed on glow-discharged holey carbon grids (Quantifoil Au R1.2/1.3). The grids were blotted for 2.5 s or 3.0 s and flash-frozen in liquid ethane cooled by liquid nitrogen with Vitrobot (Mark IV, Thermo Scientific). The cryo-EM samples were transferred to a Titan Krios operating at 300 kV equipped with Cs corrector, Gatan K3 Summit detector and GIF Quantum energy filter. Movie stacks were automatically collected using AutoEMation ^1^, with a slit width of 20 eV on the energy filter and a defocus range from −1.2 μm to −2.2 μm in super-resolution mode at a nominal magnification of 81,000×. Each stack was exposed for 2.56 s with an exposure time of 0.08 s per frame, resulting in a total of 32 frames per stack. The total dose rate was approximately 50 e^-^/Å^2^ for each stack. The stacks were motion corrected with MotionCor2 ^2^ and binned 2-fold, resulting in a pixel size of 1.087 Å/pixel. Meanwhile, dose weighting was performed ^3^. The defocus values were estimated with Gctf ^4^.

#### Cryo-EM data processing

Particles were automatically picked using Relion 3.0.6 ^5–8^ from manually selected micrographs. After 2D classification with Relion, good particles were selected and subject to two cycle of heterogeneous refinement without symmetry using cryoSPARC ^9^. The good particles were selected and subjected to Non-uniform Refinement (beta) with C1 symmetry, resulting in the 3D reconstruction for the whole structures, which was further subject to 3D classification, 3D auto-refinement and post-processing with Relion with C2 symmetry. To further improve the map quality for interface between 3E8 and ACE2-B^0^AT1 complex, the particles were C2-symmetry expanded and re-centered at the location of the interface between 3E8-ACE2 sub-complex. The re-extracted dataset was subject to focused refinement with Relion, resulting in the 3D reconstruction of better quality on the binding interface.

The resolution was estimated with the gold-standard Fourier shell correlation 0.143 criterion ^10^ with high-resolution noise substitution ^11^. Refer to Supplementary Figures S4–S6 and Supplementary Table 1 for details of data collection and processing.

#### Cryo-EM model building and structure refinement

For model building of 3E8 bound with ACE2-B^0^AT1 complex, the atomic model of the published structure S-ECD (PDB ID: 7C2L) and ACE2-B^0^AT1 complex (PDB ID: 6M18) were used as templates, which were molecular dynamics flexible fitted (MDFF) ^12^ into the whole cryo-EM map of the complex and the focused-refined cryo-EM map of the 3E8-ACE2 sub-complex, respectively. And the fitted atomic models were further manually adjusted with Coot ^13^. Each residue was manually checked with the chemical properties taken into consideration during model building. Several segments, whose corresponding densities were invisible, were not modeled. Structural refinement was performed in Phenix ^14^ with secondary structure and geometry restraints to prevent overfitting. To monitor the potential overfitting, the model was refined against one of the two independent half maps from the gold-standard 3D refinement approach. Then, the refined model was tested against the other map. Statistics associated with data collection, 3D reconstruction and model building were summarized in supplementary Table 1.

#### “Alanine walk” studies

To verify the key residues of ACE2 on binding with 3E8, Q18, E23, Q24, F28, K31, H34 and Y83 on ACE2 were changed into alanine using the overlapped PCR method with PrimeSTAR HS DNA Polymerase (Takara) and the following primers. Plasmids with mutation were expressed in mammalian expression system. After purification by Ni-NTA affinity chromatography, proteins were checked by SDS-PAGE and ELISA.

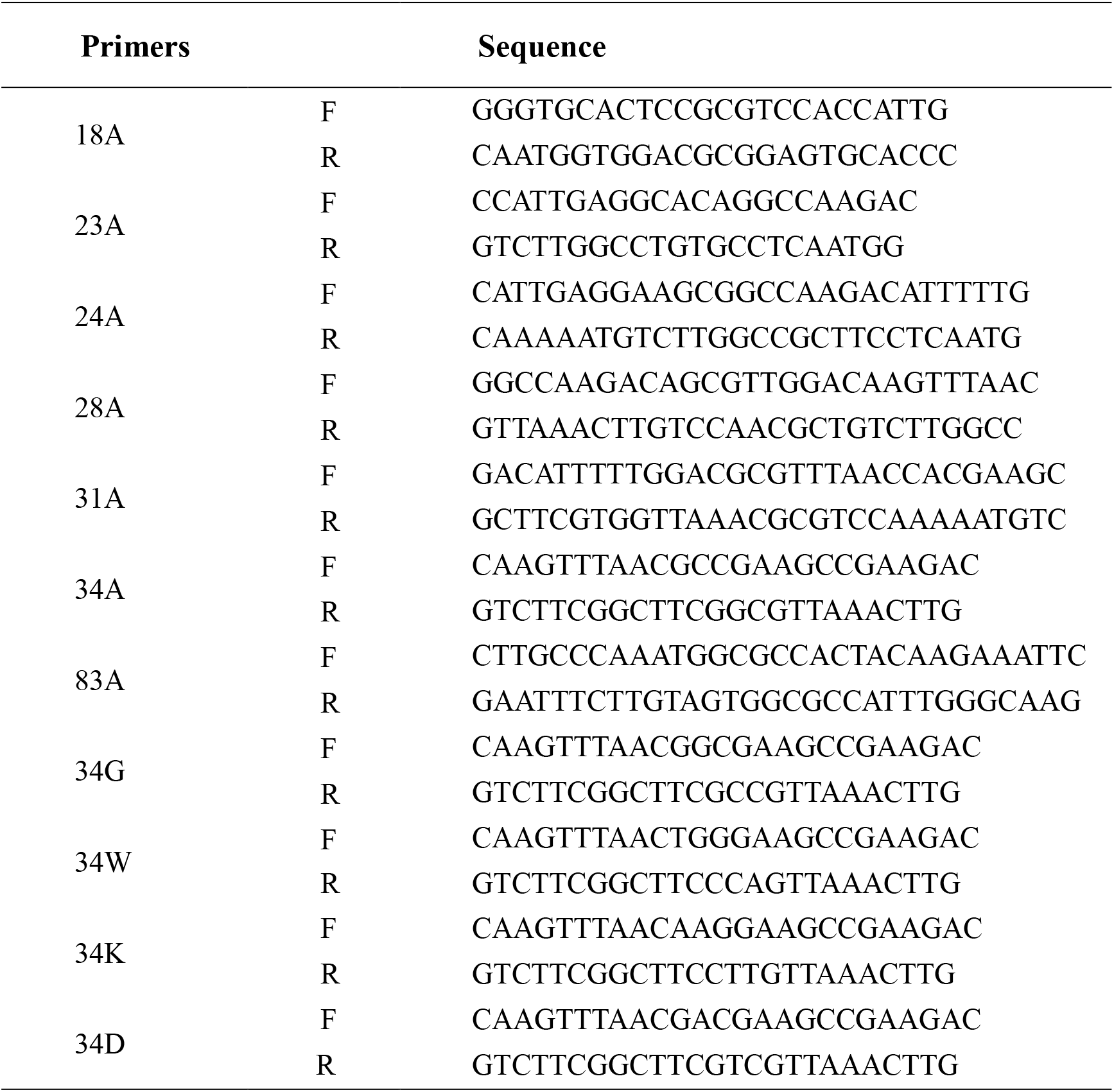

#### Toxicity studies

Five-week-old male human ACE2 “knock-in” mice on C57BL/6 background were purchased from Shanghai Model Organisms Center (Shanghai). Animal handling and procedures were approved and performed according to the requirements of the Institutional Animal Care and Use Committee (IACUC) of Shanghai Institute of Materia Medica. Five male mice were randomized into two groups: 3E8 (3 mice, 9#, 66# and 86#) and isotype (2 mice, 68# and 39#). The mice received an intravenous (i.v.) injection of 100 μl (30 mg/kg) of 3E8 or isotype. The mice were weighed and assessed for behavioral changes at 0, 24, 72 and 144 h time points after injection. Toxicities was evaluated by body weight measuring, serum biochemistry and pathology studies. After 7 days of treatment, all mice were sacrificed by cervical dislocation. Blood, hearts, livers, spleens, lungs and kidneys were collected for biochemistry and pathology studies. Organs were fixed with 10% buffered formalin and subjected to paraffin embedding before sectioned, deparaffined, rehydrated and stained with Hematoxylin and Eosin (H&E) staining.

#### Statistical analysis

Data were shown as mean ± SEM or SD. Statistical difference were calculated by Student’s t-test, with \**P* < 0.05 considered significant and *P* < 0.01 highly significant.

**Figure. S1.**
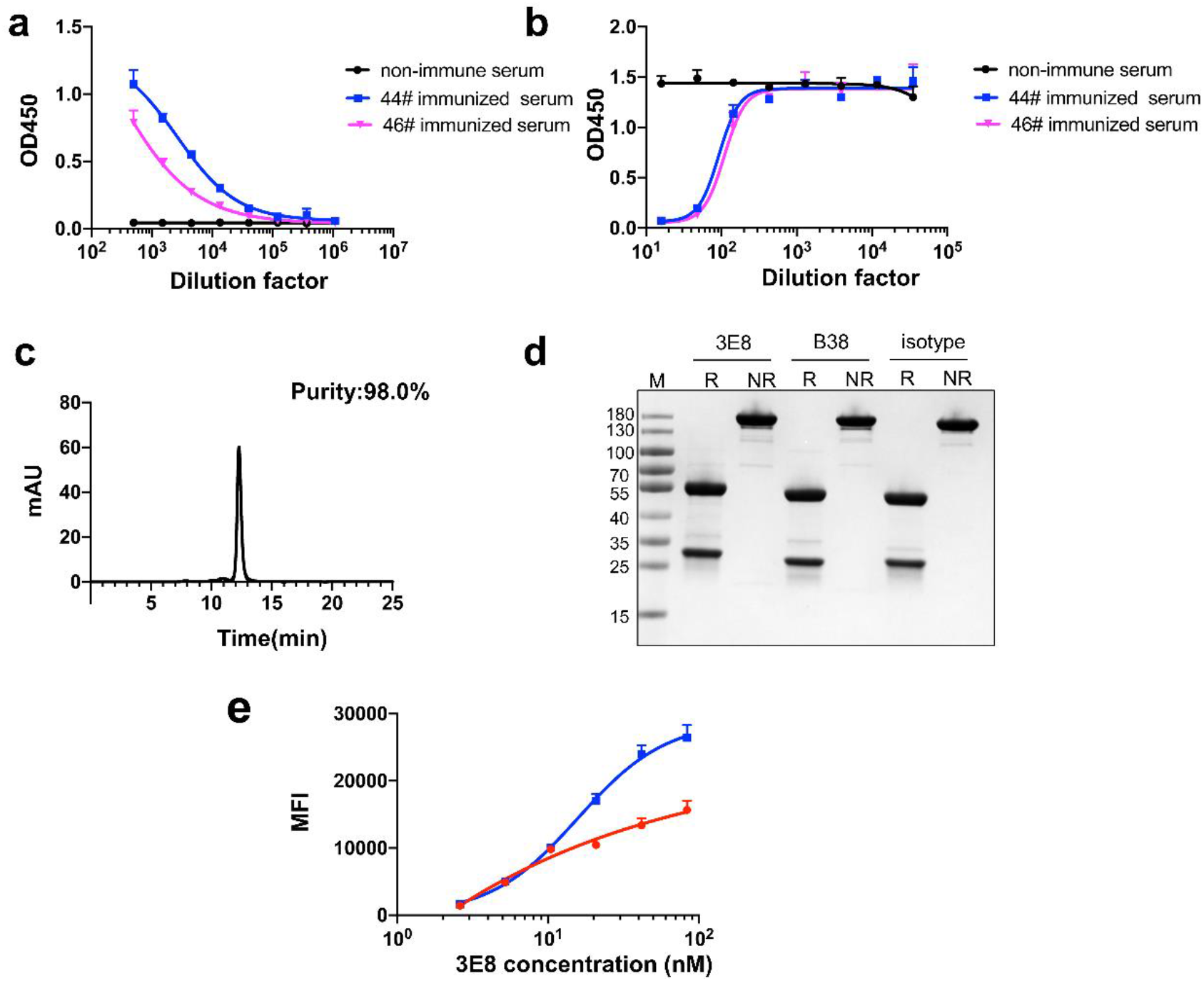
Bindings of immunized mice sera and purified 3E8 to recombinant human ACE2 protein. **a** Sera from ACE2-immunized mice bound to His-tagged recombinant human ACE2 protein. **b** Sera from ACE2-immunized mice blocked binding of SARS-CoV-2 S1-subunit to His-tagged recombinant ACE2 protein. 44# and 46# in A and B are individual immunized mice. **c** SEC profile of 3E8 on MAbPac SEC-1 column. The flow rate was 0.2 ml/min, the mobile phase was PBS buffer, and the monomer retention time was 12.28 min. **d** SDS-PAGE gels of the 3E8. R: reduced; NR: non-reduced. M: marker **e** Bindings of 3E8 to Vero E6 and HEK293 cells expressing human ACE2 measured as analyzed by flow cytometry.

**Figure. S2.**
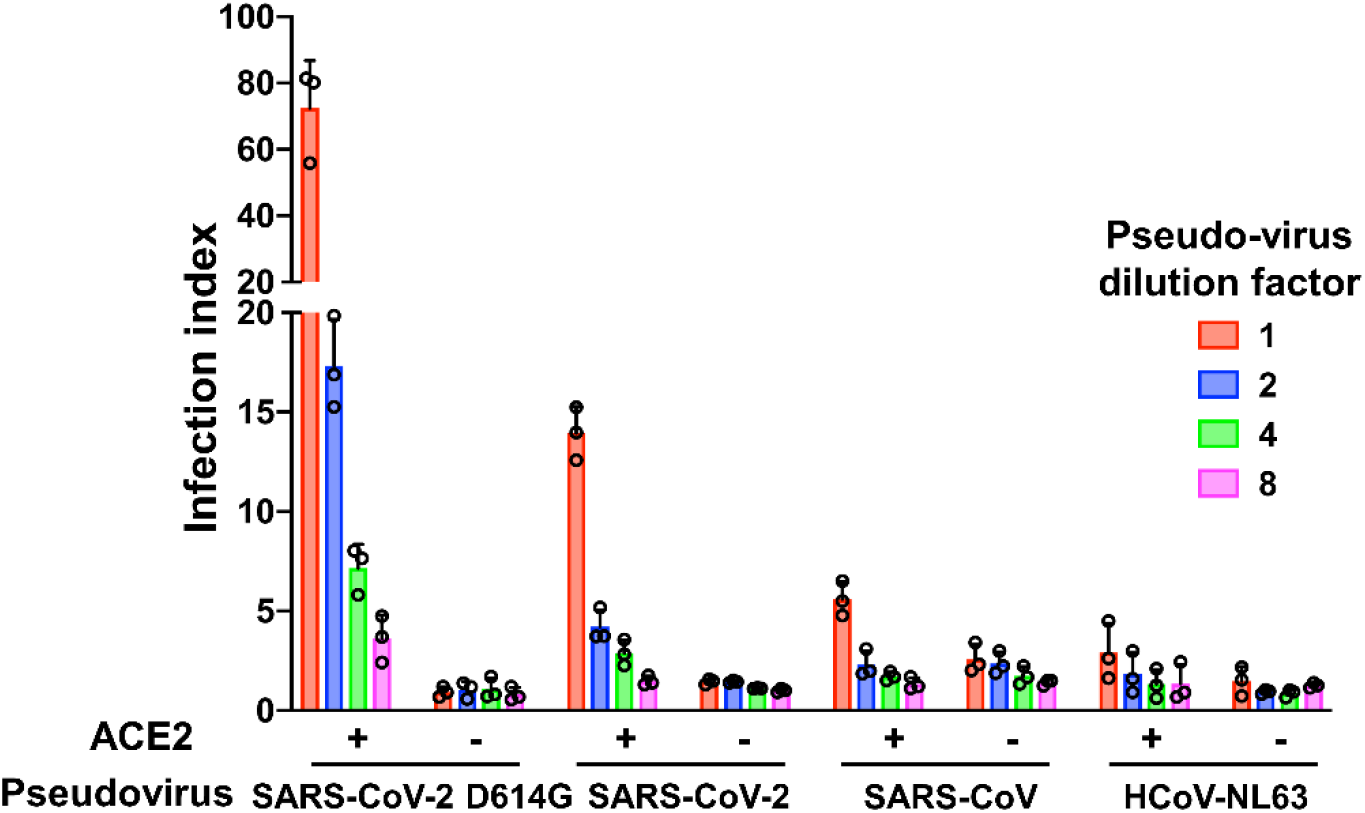
Infection of HEK293F/ACE2/EGFP cells by different pseudo-typed coronaviruses. Pseudo-typed SARS-CoV-2-D614G, SARS-CoV-2 (D614), SARS-CoV and HCoV-NL63 were constructed and infected HEK293 cells overexpressing human ACE2.

**Figure. S3.**
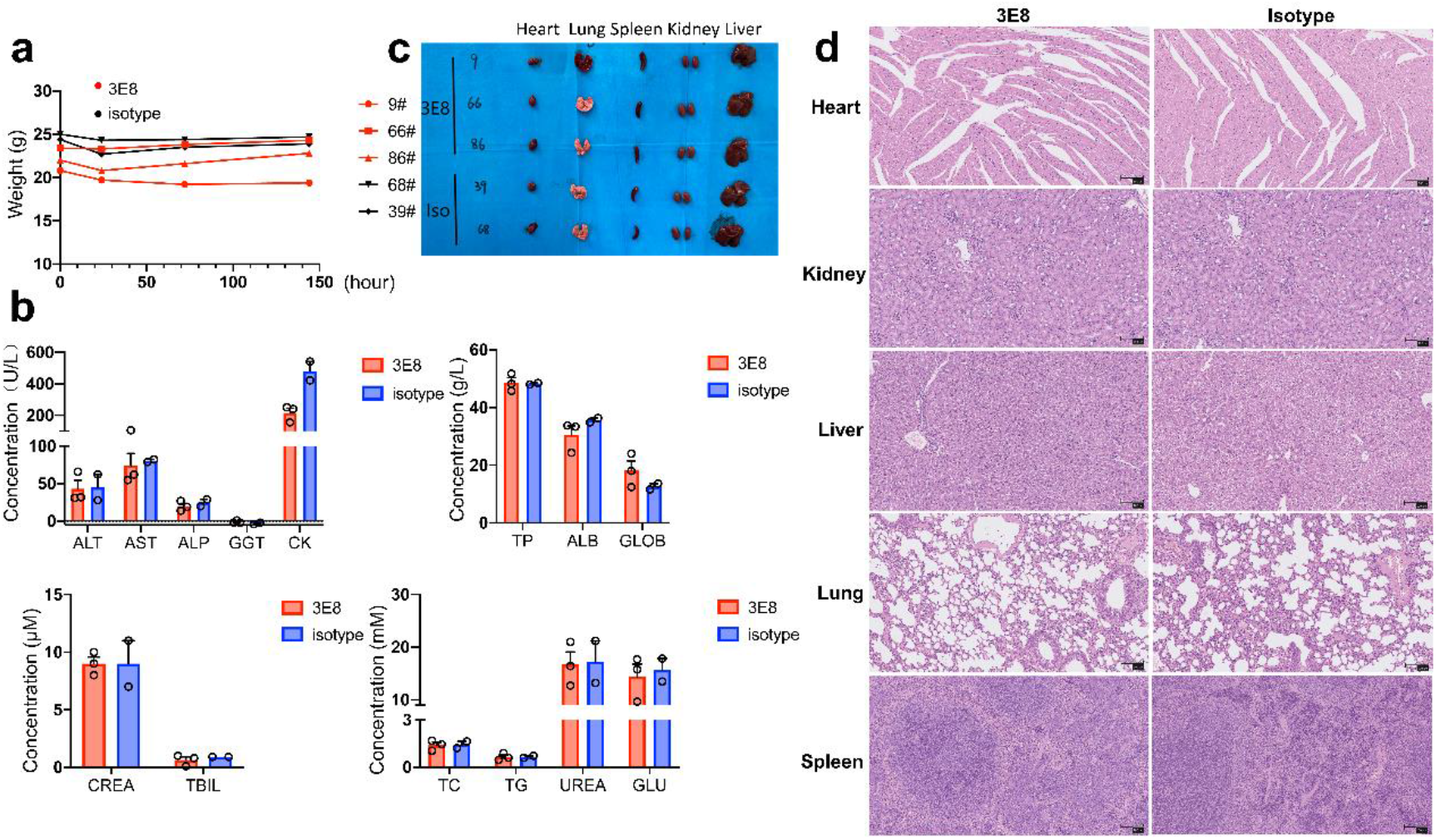
Toxicity studies of antibody 3E8 in human ACE2 “knock-in” mice. **a** The body weights of the treated mice were measured at 0, 24, 72 and 144 h time points. **b** Blood biochemistry analysis showed the plasma concentrations of alanine transaminase (ALT), aspartate transaminase (AST), alkaline phosphatase (ALP), gamma-glutamyl transpeptidase (GGT), creatine kinase (CK), total protein (TP), albumin (ALB), globulin (GLOB), creatinine (CREA), total bilirubin (TBIL), total cholesterol (TC), triglycerides (TG), urea nitrogen (UREA) and glucose (GLU). The indices of GGT and TBIL in 3E8 group and isotype group were below the normal range, but lowered levels are not signs of toxicity. **c** Shapes and sizes of major organs including hearts, livers, kidneys, spleens and lungs from mice 7 days post treatment. The organs were dissected out after the mice were sacrificed by cervical dislocation and then washed with PBS to clean out the blood. Blood clogging occurred in the lung of mouse 9# during the dissection process, which was determined technical. The mice were otherwise normal. **d** H&E staining of hearts, livers, kidneys, spleens and lungs of treated mice. No obvious pathological changes were observed. The scale of ratio above represents 100 μm.

**Figure. S4.**
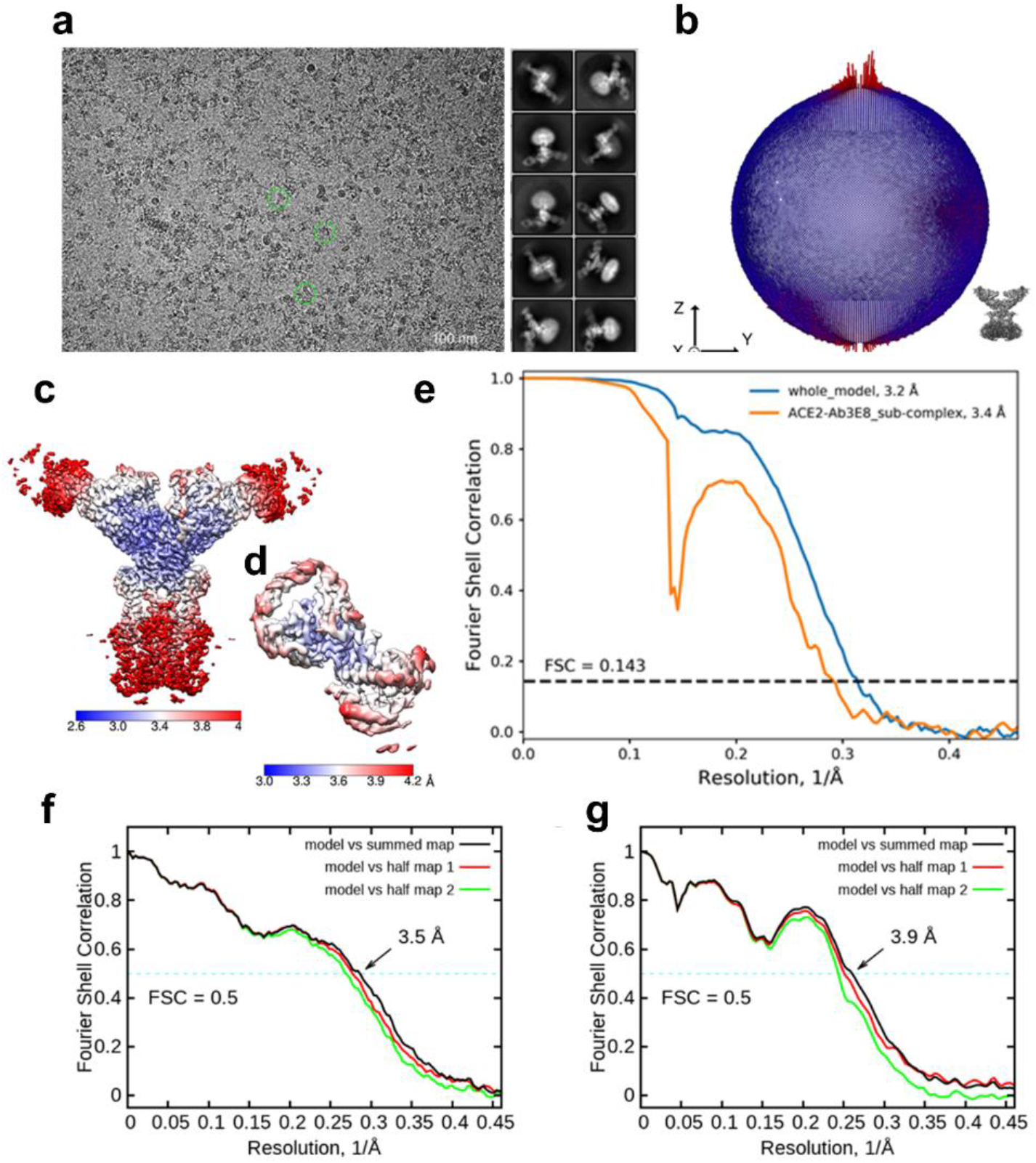
Cryo-EM analysis of 3E8 bound with ACE2-B^0^AT1 complex. **a** Representative cryo-EM micrograph and 2D class averages of cryo-EM particle images. The scale bar in 2D class averages is 10 nm. **b** Euler angle distribution in the final 3D reconstruction of 3E8 bound with ACE2-B^0^AT1 complex. **c** and **d** Local resolution maps for the 3D reconstruction of overall structure and the 3E8-ACE2 sub-complex, respectively. **e** FSC curve of the overall structure (blue) and 3E8-ACE2 sub-complex (orange). **f** FSC curve of the refined model of 3E8 bound with ACE2-B^0^AT1 complex versus the overall structure that it is refined against (black); of the model refined against the first half map versus the same map (red); and of the model refined against the first half map versus the second half map (green). The small difference between the red and green curves indicates that the refinement of the atomic coordinates is not enough overfitting. **g** FSC curve of the refined model of the 3E8-ACE2 sub-complex, which is the same as the **f**.

**Figure. S5.**
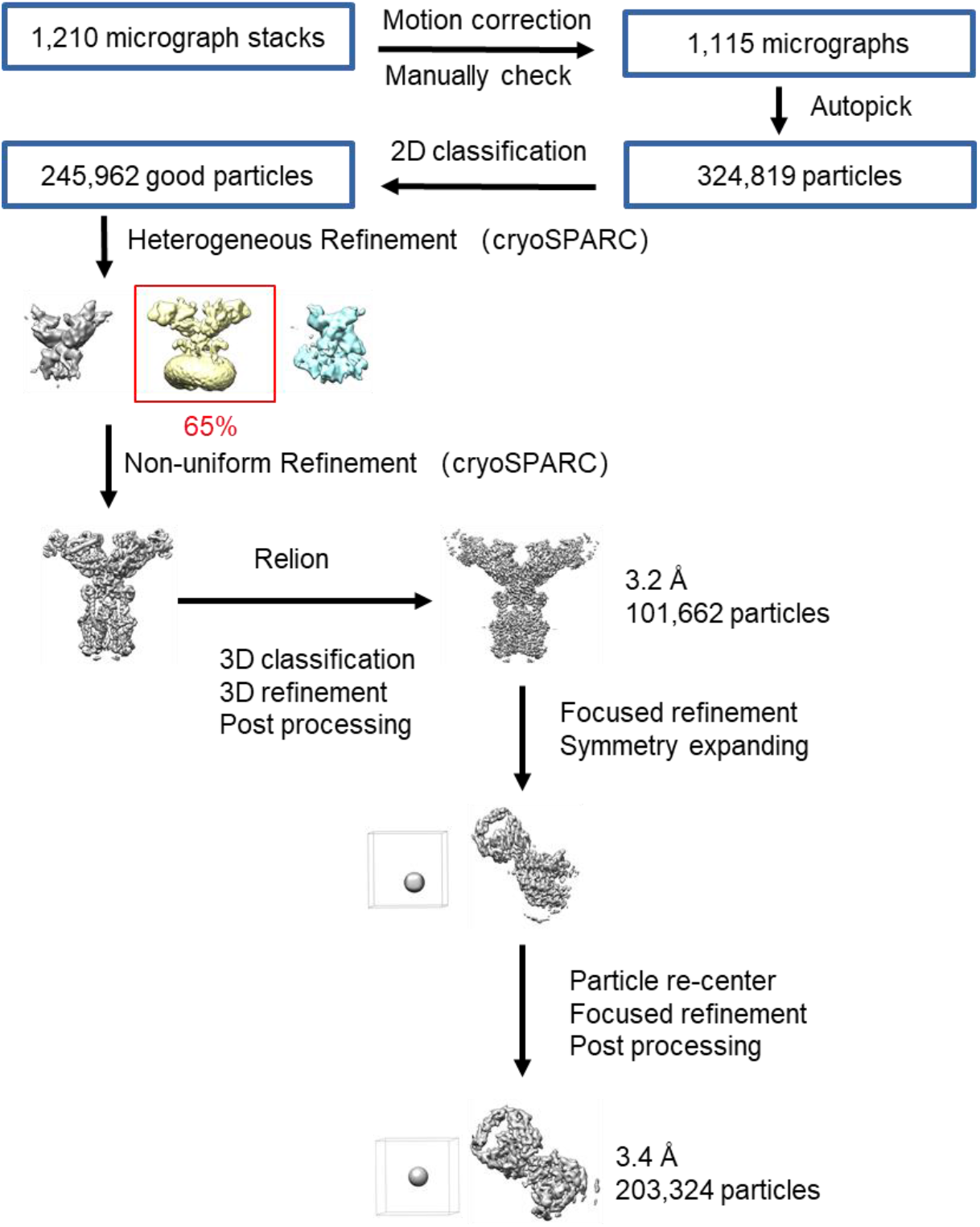
Flowchart for cryo-EM data processing. Please refer to the ‘Data Processing’ section in Methods for details.

**Figure. S6.**
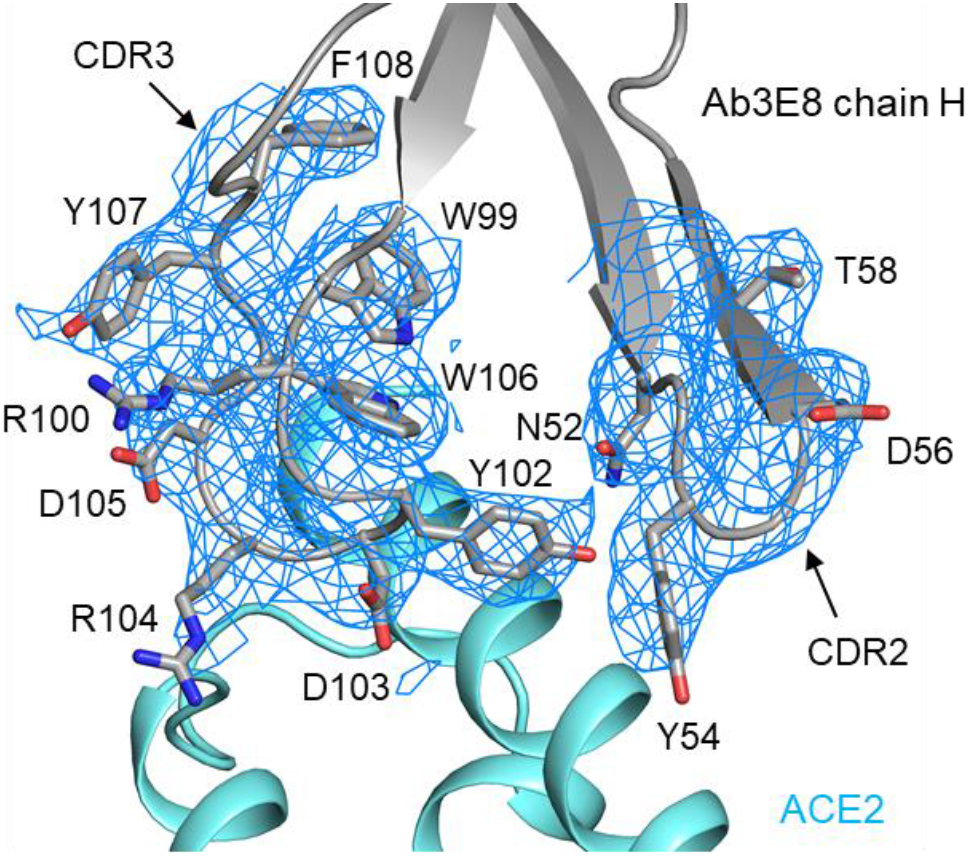
Representative cryo-EM map densities. Cryo-EM density of the interface between ACE2 and chain H of 4A8. The density is contoured at 10 σ.

**Table S1.**
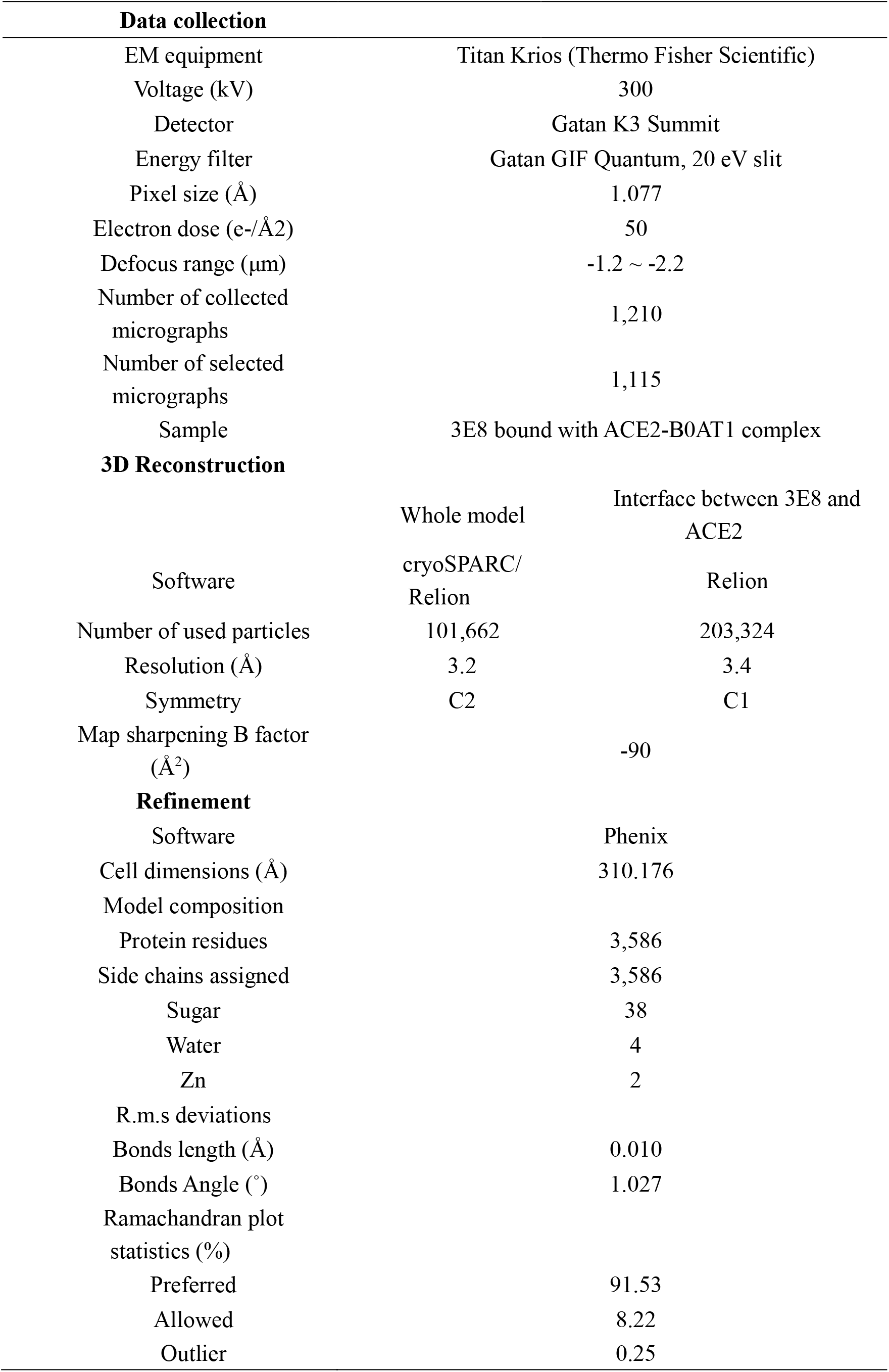
Cryo-EM data collection and refinement statistics.

## Notes

### Summary of Updates

The evolution of coronaviruses, such as SARS-CoV-2, makes broad-spectrum coronavirus preventional or therapeutical strategies highly sought after. Here we report a human angiotensin-converting enzyme 2 (ACE2)-targeting monoclonal antibody, 3E8, blocked the S1-subunits and pseudo-typed virus constructs from multiple coronaviruses including SARS-CoV-2, SARS-CoV-2 mutant variants (SARS-CoV-2-D614G, B.1.1.7, B.1.351, B.1.617.1 and P.1), SARS-CoV and HCoV-NL63, without markedly affecting the physiological activities of ACE2 or causing severe toxicity in ACE2 knock-in mice. 3E8 also blocked live SARS-CoV-2 infection in vitro and in a prophylactic mouse model of COVID-19. Cryo-EM and alanine walk studies revealed the key binding residues on ACE2 interacting with the CDR3 domain of 3E8 heavy chain. Although full evaluation of safety in non-human primates is necessary before clinical development of 3E8, we provided a potentially potent and broad-spectrum management strategy against all coronaviruses that utilize ACE2 as entry receptors and disclosed an anti-coronavirus epitope on human ACE2.

